# Fast and accurate *de novo* genome assembly from long uncorrected reads

**DOI:** 10.1101/068122

**Authors:** Robert Vaser, Ivan Sović, Niranjan Nagarajan, Mile Šikić

## Abstract

The assembly of long reads from Pacific Biosciences and Oxford Nanopore Technologies typically requires resource intensive error correction and consensus generation steps to obtain high quality assemblies. We show that the error correction step can be omitted and high quality consensus sequences can be generated efficiently with a SIMD accelerated, partial order alignment based stand-alone consensus module called Racon. Based on tests with PacBio and Oxford Nanopore datasets we show that Racon coupled with Miniasm enables consensus genomes with similar or better quality than state-of-the-art methods while being an order of magnitude faster.

Racon is available open source under the MIT license at https://github.com/isovic/racon.git.

## Introduction

With the advent of long read sequencing technologies from Pacific Biosciences (PacBio) and Oxford Nanopore Technologies (ONT), the ability to produce genome assemblies with high contiguity has received a significant fillip. However, to cope with the relatively high error rates (>5%) of these technologies assembly pipelines have typically relied on resource intensive error correction (of reads) and consensus generation (from the assembly) steps (Chin et al. 2013; Loman et al. 2015). More recent methods such as Falcon (Chin et al. 2016; https://github.com/PacificBiosciences/FALCON) and Canu (https://github.com/marbl/canu) have refined this approach and have significantly improved runtimes but are still computationally demanding for large genomes (Sović et al. 2016a). Recently, Li (Li 2016) showed that long erroneous reads can be assembled without the need for a time-consuming error-correction step. The resulting assembler, Miniasm, is an order of magnitude faster than other long-read assemblers, but produces sequences which can have >10 times as many errors as other methods (Sović et al. 2016a). As fast and accurate long-read assemblers can enable a range of applications, from more routine assembly of mammalian and plant genomes, to structural variation detection, improved metagenomic classification and even online, “read until” assembly (Loose et al. 2016), a fast and accurate consensus module is a critical need. This was also noted by Li (Li 2016), highlighting that fast assembly was only feasible if a consensus module matching the speed of Minimap and Miniasm was developed as well.

Here we address this need by providing a very fast consensus module called Racon (for **Ra**pid **Con**sensus), which when paired with a fast assembler such as Miniasm, enables the efficient construction of genome sequences with high accuracy (Q30) even without an error correction step. Assemblies from this pipeline (Miniasm+Racon) are comparable to those from state-of-the-art methods such as Falcon and Canu, while being an order of magnitude faster in many cases. Racon provides a first standalone, platform-independent consensus module for long and erroneous reads and can also be used as a fast and accurate read correction tool.

## Results

Racon is designed as a user friendly standalone consensus module that is not explicitly tied to any *de novo* assembly method or sequencing technology. It reads multiple input formats (GFA, FASTA, FASTQ, SAM, MHAP and PAF), allowing simple interoperability and modular design of new pipelines. Even though other stand-alone consensus modules, such as Quiver (Chin et al. 2013) and Nanopolish (Loman et al. 2015) exist, they require sequencer specific input and are intended to be applied *after* the consensus phase of assembly to further polish the sequence. Racon is run with sequencer-independent input, is robust enough to work with uncorrected read data and is designed to rapidly generate high-quality consensus sequences. These sequences can be further polished with Quiver or Nanopolish or by applying Racon for more iterations.

Racon can take as input a set of raw backbone sequences, a set of reads and a set of overlaps between reads and backbone sequences. Overlaps can be generated using any overlapper which supports either the MHAP or PAF output formats, such as Minimap (Li 2016), MHAP (Berlin et al. 2015) or GraphMap (Sović et al. 2016b). In our tests, we used Minimap as the overlapper as it was the fastest and provided reasonable results. Racon uses the overlap information to construct a partial order alignment graph, using a Single Instruction Multiple Data (SIMD) implementation to accelerate the process (SPOA). More details on Racon and SPOA can be found in the Methods section.

For the purpose of evaluation, we paired Racon with Miniasm to form a fast and accurate *de novo* assembly pipeline (referred to here as *Miniasm+Racon*), which we then compared to other state-of-the-art *de novo* assembly tools for third generation sequencing data (i.e. Falcon and Canu). Note that Falcon and Canu have previously been benchmarked with other assembly methods such as PBcR and a pipeline from Loman *et al.* (Loman et al. 2015) and shown to produce high quality assemblies with improved running times (Sović et al. 2016a). Assembly pipelines were evaluated in terms of consensus sequence quality (**Table 1**), runtime and memory usage (**Table 2**; **Figure 1**), and scalability with respect to genome size (**Figure 2**), on several PacBio and Oxford Nanopore datasets (see Methods).

**Figure 1.**
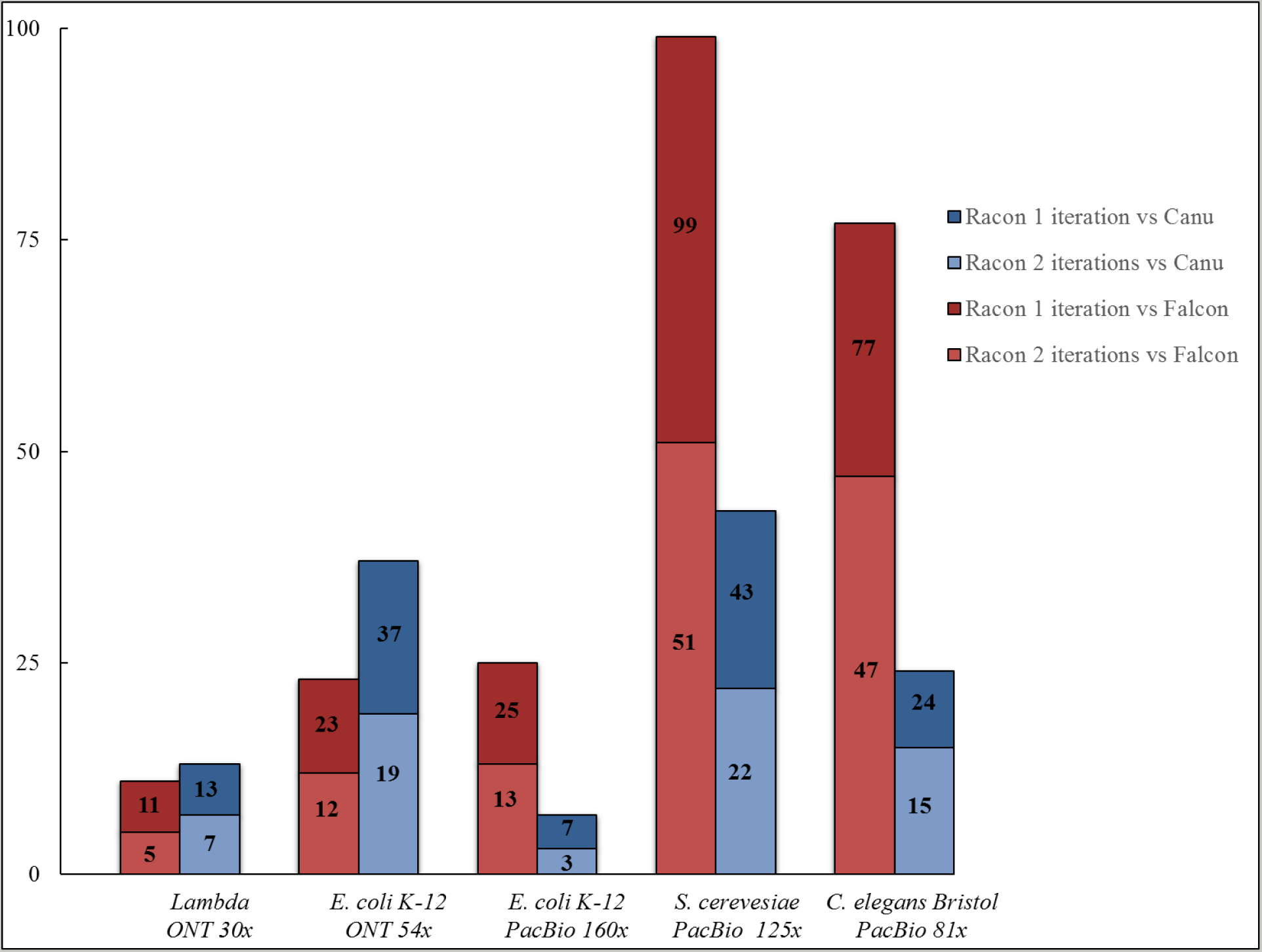
Racon’s speed-up when compared to Falcon and Canu.

**Figure 2.**
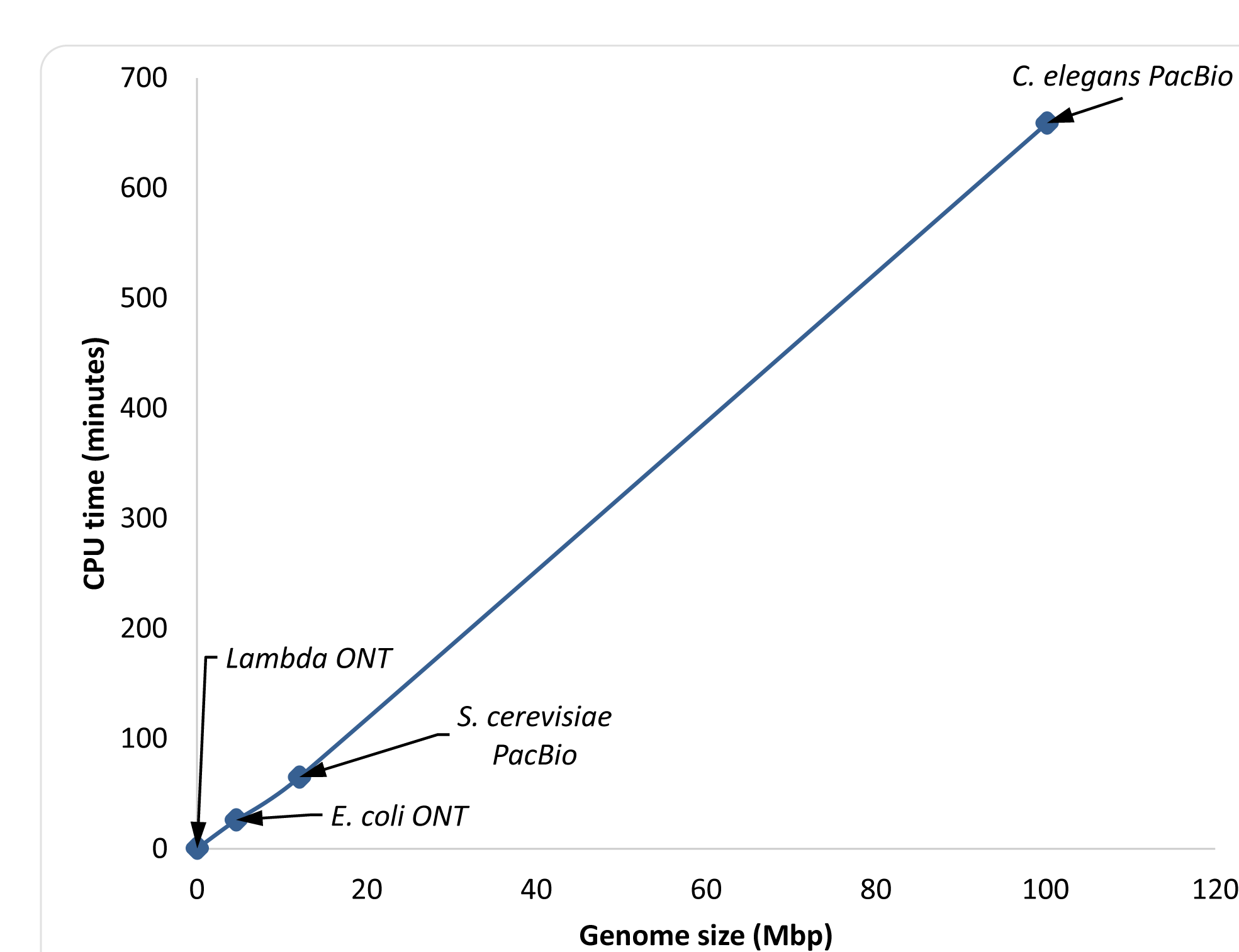
Scalability of Racon as a function of genome size. Read coverage was subsampled to be 81× (limited by the *C. elegans* dataset) and the figure shows results for one iteration of Racon.

**Table 1.** Assembly and consensus results accross 5 datasets of varying genome length and sequencing data type.

|  |  | Miniasm+Racon<br>1 iteration | Miniasm+Racon<br>2 iterations | Canu | Falcon |
| --- | --- | --- | --- | --- | --- |
| *Lambda*<br>ONT<br>30× | Ref. genome size [bp] | 48502 | 48502 | 48502 | 48502 |
|  | Total bases [bp] | 47917 | 47874 | 25077 | 7212 |
|  | Ref. chromosomes [#] | 1 | 1 | 1 | 1 |
|  | Contigs [#] | 1 | 1 | 1 | 1 |
|  | Aln. bases ref. [bp] | 48438 (99.87%) | 48425 (99.84%) | 25833 (53.26%) | 7483 (15.43%) |
|  | Aln. bases query [bp] | 47917 (100.00%) | 47874 (100.00%) | 25077 (100.00%) | 7212 (100.00%) |
|  | Avg. Identity | 97.57 | 97.90 | 96.87 | 95.77 |
|  | CPU time [min] | 0.22 | 0.43 | 2.87 | 2.30 |
|  | Memory [GB] | 0.066 | 0.066 | 1.897 | 0.854 |
| *E. coli*<br>K-12<br>ONT<br>R7.3<br>54× | Ref. genome size [bp] | 4641652 | 4641652 | 4641652 | 4641652 |
|  | Total bases [bp] | 4637221 | 4632092 | 4601503 | 4580230 |
|  | Ref. chromosomes [#] | 1 | 1 | 1 | 1 |
|  | Contigs [#] | 1 | 1 | 1 | 1 |
|  | Aln. bases ref. [bp] | 4640867 (99.98%) | 4641323 (99.99%) | 4631173 (99.77%) | 4627613 (99.70%) |
|  | Aln. bases query [bp] | 4636904 (99.99%) | 4632089 (100.00%) | 4601365 (100.00%) | 4580230 (100.00%) |
|  | Avg. Identity | 99.13 | 99.32 | 99.28 | 98.84 |
|  | CPU time [min] | 36 | 70 | 1328 | 829 |
|  | Memory [GB] | 3.32 | 3.32 | 4.03 | 12.29 |
| *E. coli*<br>K-12<br>PacBio<br>P6C4<br>160× | Ref. genome size [bp] | 4641652 | 4641652 | 4641652 | 4641652 |
|  | Total bases [bp] | 4653227 | 4645508 | 4664416 | 4666788 |
|  | Ref. chromosomes [#] | 1 | 1 | 1 | 1 |
|  | Contigs [#] | 1 | 1 | 1 | 1 |
|  | Aln. bases ref. [bp] | 4641501 (100.00%) | 4641439 (100.00%) | 4641652 (100.00%) | 4641652 (100.00%) |
|  | Aln. bases query [bp] | 4653139 (100.00%) | 4645508 (100.00%) | 4664416 (100.00%) | 4666788 (100.00%) |
|  | Avg. Identity | 99.63 | 99.90 | 99.99 | 99.90 |
|  | CPU time [min] | 116 | 225 | 773 | 2908 |
|  | Memory [GB] | 9.74 | 9.74 | 3.59 | 9.93 |
| *S. cerevisiae*<br>W303<br>PacBio<br>P4C2<br>127× | Ref. genome size [bp] | 12071326 | 12071326 | 12071326 | 12071326 |
|  | Total bases [bp] | 12071319 | 12051772 | 12402332 | 12003077 |
|  | Ref. chromosomes [#] | 16 | 16 | 16 | 16 |
|  | Contigs [#] | 30 | 30 | 29 | 44 |
|  | Aln. bases ref. [bp] | 11939290 (98.91%) | 11939845 (98.91%) | 12042102 (99.76%) | 11922591 (98.77%) |
|  | Aln. bases query [bp] | 11962050 (99.09%) | 11942005 (99.09%) | 12269365 (98.93%) | 11900584 (99.15%) |
|  | Avg. Identity | 99.44 | 99.73 | 99.79 | 99.58 |
|  | CPU time [min] | 150 | 290 | 6375 | 14808 |
|  | Memory [GB] | 16.07 | 16.07 | 3.65 | 4.78 |
| *C. elegans*<br>PacBio<br>P6C4<br>81× | Ref. genome size [bp] | 100272607 | 100272607 | 100272607 | 100272607 |
|  | Total bases [bp] | 106352656 | 106387537 | 106687886 | 105858394 |
|  | Ref. chromosomes [#] | 6 | 6 | 6 | 6 |
|  | Contigs [#] | 77 | 77 | 134 | 242 |
|  | Aln. bases ref. [bp] | 100017755 (99.75%) | 100015191 (99.74%) | 100166301 (99.89%) | 99295695 (99.03%) |
|  | Aln. bases query [bp] | 101710096 (95.63%) | 101772785 (95.66%) | 102928910 (96.48%) | 102008289 (96.36%) |
|  | Avg. Identity | 99.44 | 99.74 | 99.89 | 99.74 |
|  | CPU time [min] | 1561 | 2567 | 37852 | 119766 |
|  | Memory [GB] | 85.53 | 85.53 | 10.16 | 7.59 |

**Table 2.** Resource usage for various parts of the Miniasm+Racon assembly pipeline. Results are presented in the format “CPU time [s] / Maximum memory [GB]”.

|  | *Lambda* ONT | *E. coli* ONT | *E. coli* PacBio | *S. cerevisiae* PacBio | *C. elegans* PacBio |
| --- | --- | --- | --- | --- | --- |
| Minimap overlap | 0.58 / 0.038 | 170 / 3 | 670 / 10 | 1393 / 16 | 33203 / 48 |
| Miniasm | 0.01 / 0.001 | 4 / 0.06 | 25 / 0.39 | 31 / 0.46 | 236 / 3 |
| Minimap mapping 1st iter. | 0.07 / 0.007 | 14 / 0.23 | 37 / 0.23 | 86 / 0.26 | 814 / 1 |
| Racon consensus 1st iter. | 13 / 0.066 | 1995 / 3 | 6216 / 8 | 7470 / 14 | 59393 / 86 |
| Minimap mapping 2nd iter. | 0.08 / 0.005 | 16 / 0.23 | 43 / 0.23 | 97 / 0.26 | 880 / 1 |
| Racon consensus 2nd iter. | 12 / 0.06 | 1976 / 2 | 6537 / 6 | 8338 / 13 | 59493 / 71 |
| Total CPU time / Max. mem. | 26 / 0.066 | 4175 / 3 | 13528 / 10 | 17415 / 16 | 154019 / 86 |

As can be seen from **Table 1**, all assembly pipelines were able to produce assemblies with high coverage of the reference genome and in a few contigs. Canu, Falcon and the Miniasm+Racon pipeline also constructed sequences with comparable sequence identity to the reference genome, with the iterative use of Racon serving as a polishing step for obtaining higher sequence identity. In addition, the Miniasm+Racon pipeline was found to be significantly faster for all datasets, with a 3-23× speedup compared to Canu and 7-51× speedup compared to FALCON (with two Racon iterations; **Figure 1**).

Racon's speedup was more pronounced for larger genomes and is likely explained by the observation that it scales linearly with genome size (for fixed coverage; **Figure 2**).

The runtime of the Miniasm+Racon pipeline was dominated by the time for the consensus generation step in Racon, highlighting that this step is still the most compute intensive one for small genomes (**Table 2**). However, the results in **Table 2** suggest that for larger genomes the overlap computation stage can catch up in terms of resource usage. Furthermore, if a polishing stage is used, this would typically be more resource intensive. Comparison of the results of the various assembly pipelines after a polishing stage confirmed that the use of Racon provided better results than just the Miniasm assembly (avg. identity of 99.80% vs 98.06%) and that the Miniasm+Racon assembly matched the best reported sequence quality for this dataset (from the Loman *et al.* pipeline; Sović et al. 2016a), while providing a better match to the *actual size* of the reference genome (4641652 bp; **Table 3**). We additionally observed that Nanopolish executed >6× faster on Miniasm+Racon contigs than on raw Miniasm assemblies (248.28 CPUh vs. 1561.80 CPUh), and the Miniasm+Racon+Nanopolish approach achieved the same sequence quality as the original Loman *et al.* pipeline, while being much faster.

**Table 3.**
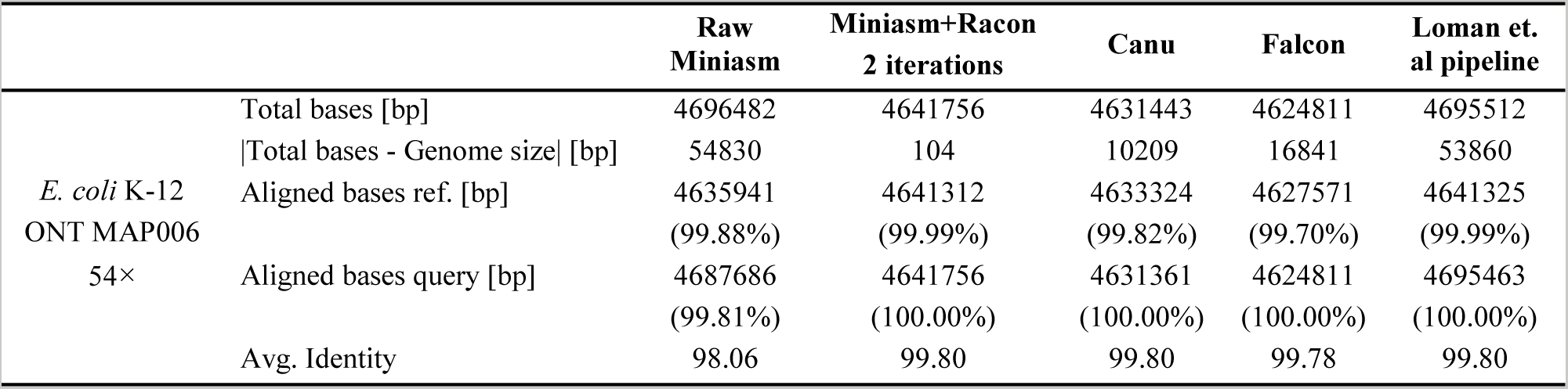
Results after polishing assemblies with Nanopolish.

Finally, we also evaluated Racon's use as an error-correction module. We noted that Racon corrected reads had error rates comparable to Falcon and Canu but provided better coverage of the genome (**Table 4**). Overall, Nanocorrect (Loman et al. 2015) had the best results in terms of error rate but it had lower reference coverage and was more than two orders of magnitude slower than Racon.

**Table 4.**
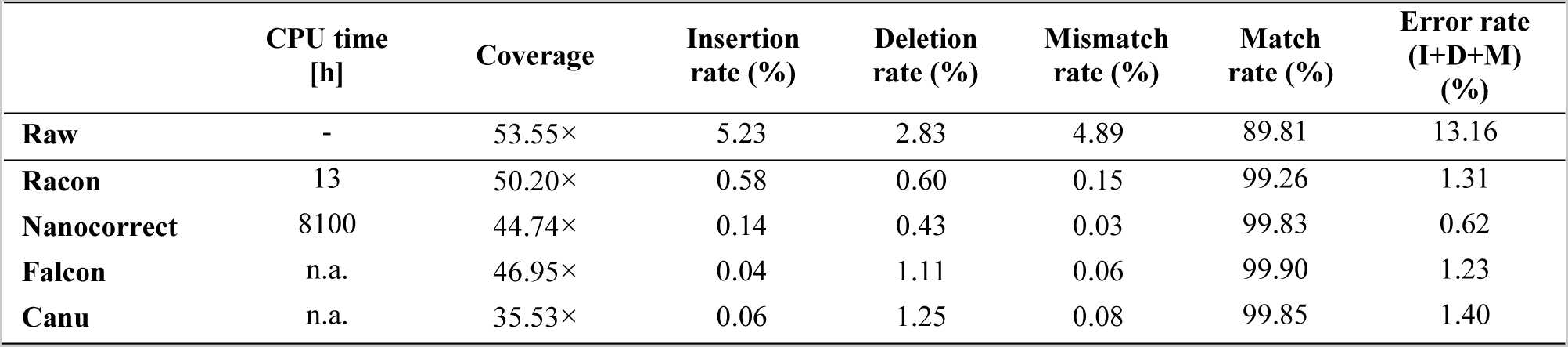
Comparison of error-correction modules on *E. coli* K-12 MAP006 R7.3 54× dataset. Values presented in the table are median values of the error and match rate estimates.

|  | CPU time<br>[h] | Coverage | Insertion<br>rate (%) | Deletion<br>rate (%) | Mismatch<br>rate (%) | Match<br>rate (%) | Error rate<br>(I+D+M)<br>(%) |
| --- | --- | --- | --- | --- | --- | --- | --- |
| **Raw** | - | 53.55× | 5.23 | 2.83 | 4.89 | 89.81 | 13.16 |
| **Racon** | 13 | 50.20× | 0.58 | 0.60 | 0.15 | 99.26 | 1.31 |
| **Nanocorrect** | 8100 | 44.74× | 0.14 | 0.43 | 0.03 | 99.83 | 0.62 |
| **Falcon** | n.a. | 46.95× | 0.04 | 1.11 | 0.06 | 99.90 | 1.23 |
| **Canu** | n.a. | 35.53× | 0.06 | 1.25 | 0.08 | 99.85 | 1.40 |

## Discussion

The principal contribution of this work is to take the concept of fast, error-correction-free, long read assembly, as embodied by the recently developed program Miniasm, to its logical end. Miniasm is remarkably efficient and effective in taking erroneous long reads and producing contig sequences that are structurally accurate (Sović et al. 2016a). However, assemblies from Miniasm do not match up in terms of sequence quality when compared with the best assemblies that can be produced with existing assemblers. This serves as a significant barrier for adopting this ‘light-weight’ approach to assembly, despite its attractiveness for greater adoption of *de novo* assembly methods in genomics. In this work we show that the sequence quality of a correction-free assembler can indeed be efficiently boosted to a quality comparable to other resource-intensive state-of-the-art assemblers. This makes the tradeoff offered much more attractive and the concept of a correction-free assembler more practically useful.

Racon is able to start from uncorrected contigs and raw reads and still generate accurate sequences efficiently because it exploits the development of a Single Instruction Multiple Data (SIMD) version of the robust partial order alignment framework. This makes the approach scalable to large genomes and general enough to work with data from very different sequencing technologies. With the increasing interest in the development of better third generation assembly pipelines, we believe that Racon can serve as useful plug-in consensus module that enables software reuse and modular design.

## Methods

Racon is based on the *Partial Order Alignment* (POA) graph approach (Lee et al. 2002; Lee 2003) and we report the development of a Single Instruction Multiple Data (SIMD) version that significantly accelerates this analysis. An overview of Racon’s steps is given in **Figure 3**. The entire process is also shown in detail in **Algorithm 1**.

**Figure 3.**
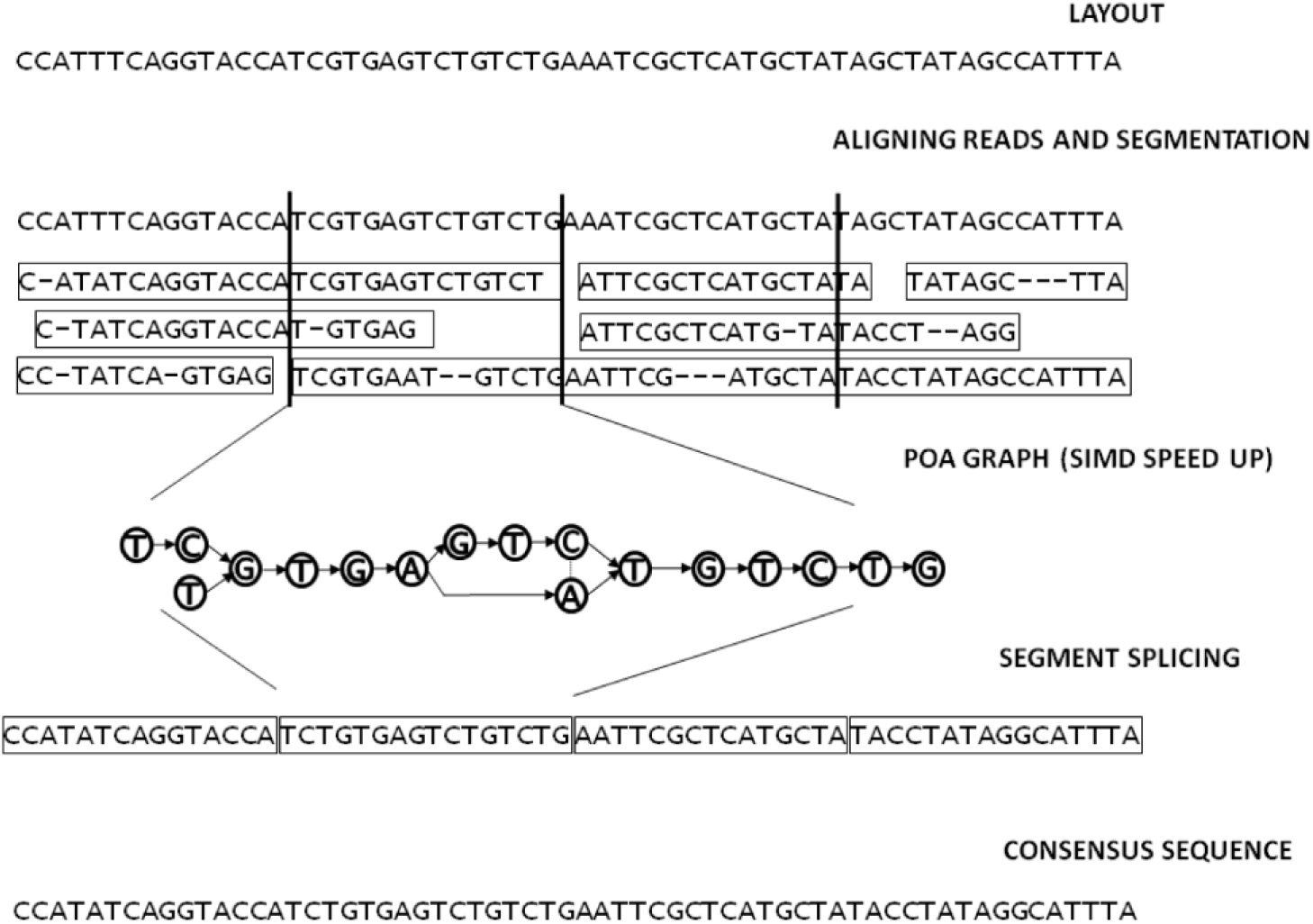
Overview of the Racon consensus process.

To perform consensus calling (or error-correction), Racon depends on an input set of query-to-target overlaps (query is the set of reads, while a target is either a set of contigs in the consensus context, or a set of reads in the error-correction context). Racon then loads the overlaps and performs simple filtering (**Algorithm 1**, lines 1−3; **Algorithm 2**): (I) at most one overlap per read is kept in consensus context (in error-correction context this particular filtering is disabled), and (II) overlaps which have high error-rate (i.e. |1 − min(*d_q_*, *d_t_*)/max(*d_q_*,*d_t_*)| ≥ *e*, where *d_q_* and *d_t_* are the lengths of the overlap in the query and the target respectively and *e* is a user-specified error-rate threshold) are removed. For each overlap which survived the filtering process, a fast edit-distance based alignment is performed (Myers 1999) (**Algorithm 1**, lines 4−10). We used Edlib implementation of Myers algorithm (https://github.com/Martinsos/edlib). This alignment is needed only to split the reads into chunks which fall into particular non-overlapping windows on the backbone sequence. Each window is then processed independently in a separate thread by constructing a POA graph using SIMD acceleration and calling the consensus of the window. The final consensus sequence is then constructed by splicing the individual window consensuses together (per contig or read to be corrected).

**Algorithm 1.**
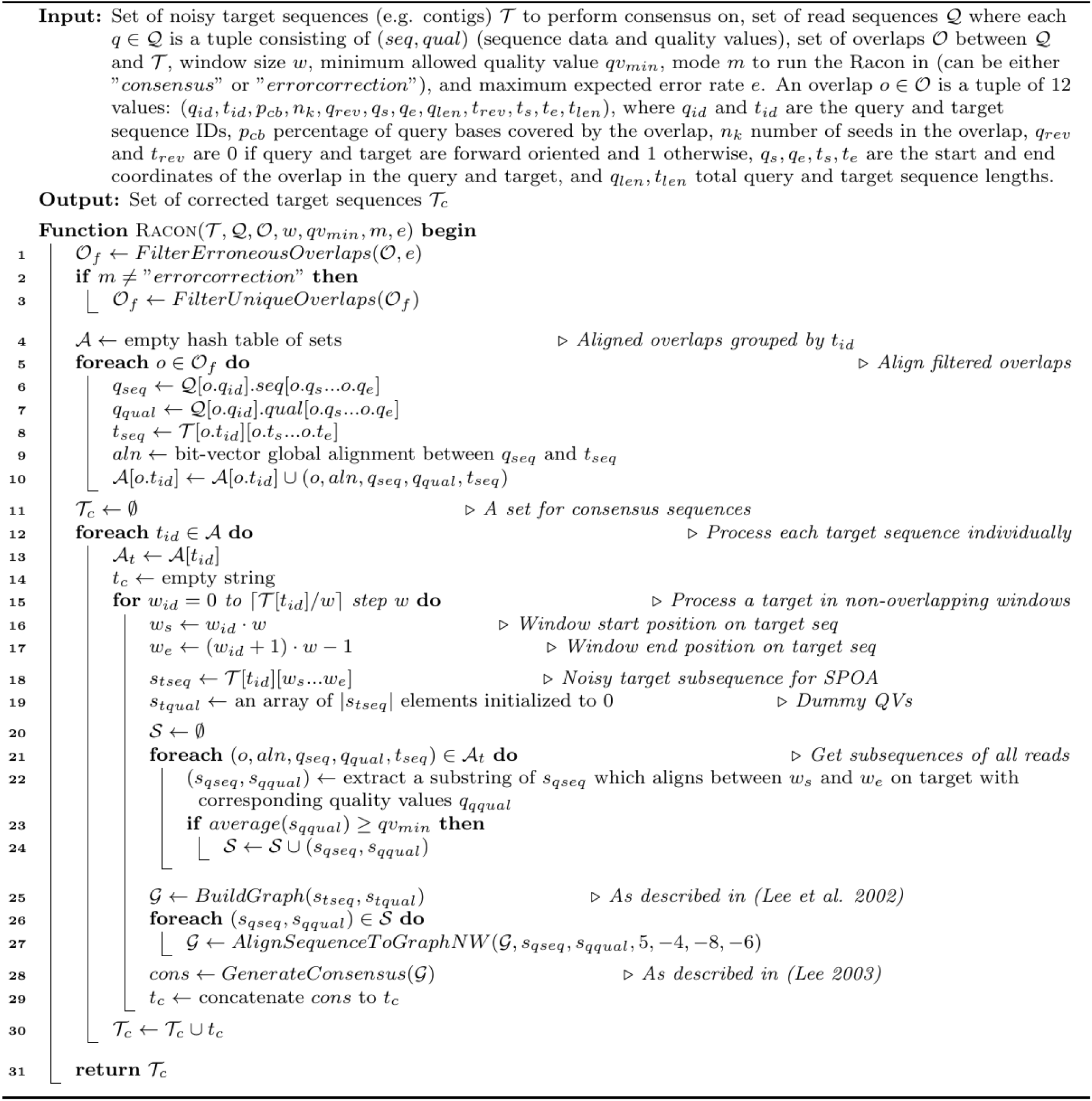
The Racon algorithm for consensus generation.

**Algorithm 2.**
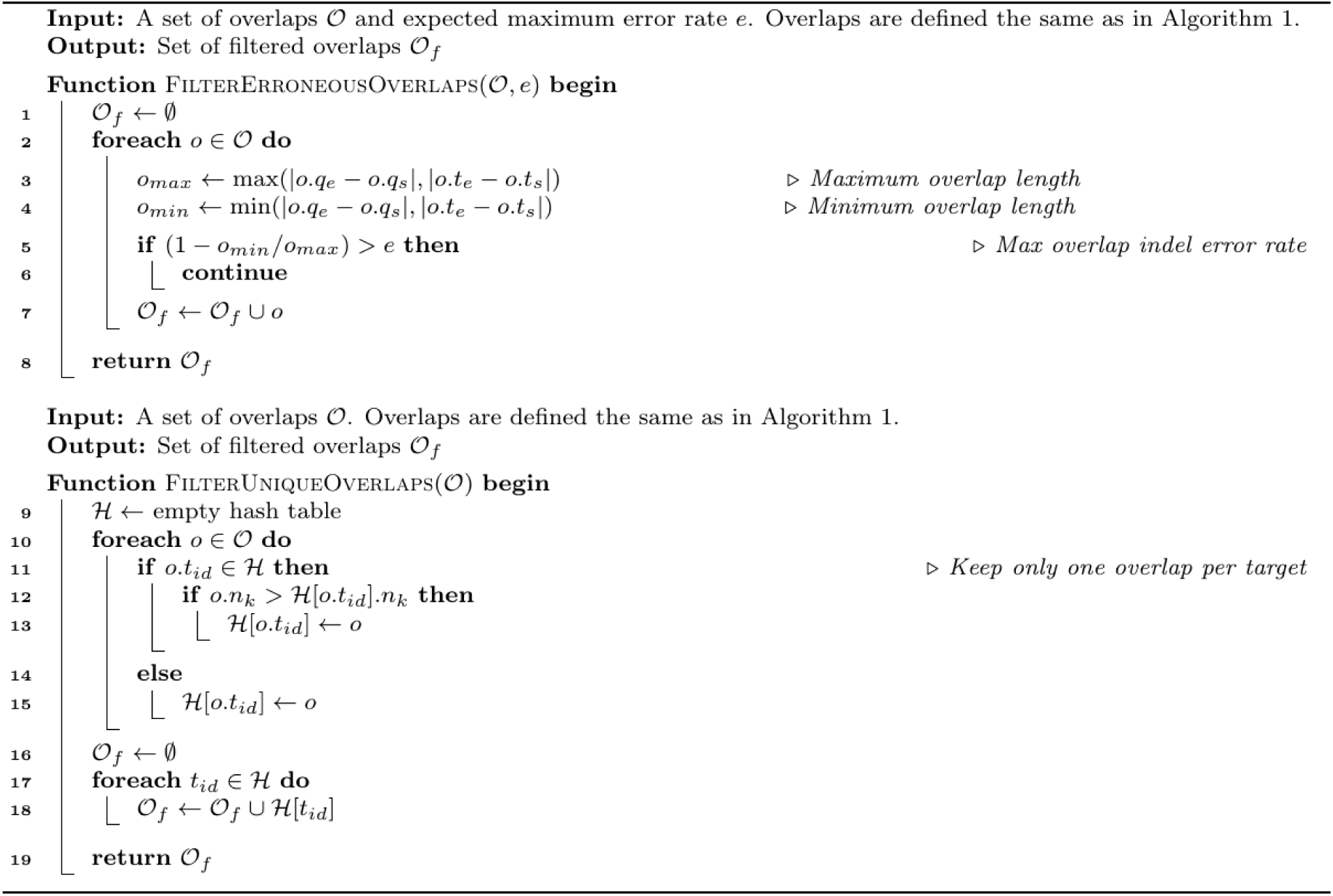
Functions for filtering overlaps in Racon.

### Partial order alignment and SIMD vectorization

POA performs Multiple Sequence Alignment (MSA) through a directed acyclic graph (DAG), where nodes are individual bases of input sequences, and weighted, directed edges represent whether two bases are neighboring in any of the sequences. Weights of the edges represent the multiplicity (coverage) of each transition. Alternatively, weights can be set according to the base qualities of sequenced data. The alignment is carried out directly through dynamic programming (DP) between a new sequence and a pre-built graph. While the regular DP for pairwise alignment has time complexity of *O*(3*nm*), where *n* and *m* are the lengths of the sequences being aligned, the sequence to graph alignment has a complexity of *O*((2*n_p_* + 1) *n*|*V*|), where *n_p_* is the average number of predecessors in the graph and |*V*| is the number of nodes in the graph (Lee et al. 2002).

Consensus sequences are obtained from a built POA graph by performing a topological sort and processing the nodes from left to right. For each node *v*, the highest-weighted in-edge *e* of weight *e_w_* is chosen, and a score is assigned to *v* such that *scores*[*v*] = *e_w_* + *scores*[*w*] where *w* is the source node of the edge *e* (Lee 2003). The node *w* is marked as a predecessor of *v*, and a final consensus is generated by performing a traceback from the highest scoring node *r*. In case *r* is an internal node (*r* has out edges), Lee (Lee 2003) proposed the idea of branch completion, where all scores for all nodes except *scores*[*r*] would be set to a negative value, and the traversal would continue from *r* as before, with the only exception that nodes with negative scores could not be added as predecessors to any other node.

One of the biggest advantages of POA compared to other MSA algorithms is its speed, with its linear time complexity in the number of sequences (Lee et al. 2002). However, even though it is faster than other MSA algorithms, the implementations of POA in current error-correction modules, such as Nanocorrect, are prohibitively slow for larger datasets. In order to increase the speed of POA while retaining its robustness, we explored a Single Instruction Multiple Data (SIMD) version of the algorithm (SPOA).

SPOA (**Figure 4**; **Algorithm 3**) is inspired by the Rognes and Seeberg Smith-Waterman intra-set parallelization approach (Rognes and Seeberg 2000). It places the SIMD vectors parallel to the query sequence (the read), while placing a graph on the other dimension of the DP matrix (**Figure 4**). In our implementation, the matrices used for tracking the maximum local-alignment scores ending in gaps are stored entirely in memory (**Algorithm 3**, line 8 and 10). These matrices are needed to access scores of predecessors of particular nodes during alignment. Unlike regular Gotoh alignment, for each row in the POA DP matrix all its predecessors (via in-edges of the corresponding node in graph) need to be processed as well (**Algorithm 3**, line 17). All columns are then processed using SIMD operations in a query-parallel manner and the values of Gotoh’s vertical matrix (**Algorithm 3**, line 20) and a partial update to Gotoh’s main scoring matrix (**Algorithm 3**, line 24) are calculated. SIMD operations in **Algorithm 3** process 8 cells of the DP matrix at a time (16-bit registers). A temporary variable is used to keep the last cell of the previous vector for every predecessor (**Algorithm 3**, lines 21−23), which is needed to compare the upper-left diagonal of the current cell to the cell one row up. Processing the matrix horizontally is not performed using SIMD operations due to data dependencies (each cell depends on the result of the cell to the left of it), and are instead processed linearly (**Algorithm 3**, lines 25−33). SPOA uses shifting and masking to calculate every particular value of a SIMD vector individually (**Algorithm 3**, lines 29-31). After the alignment is completed, the traceback is performed (**Algorithm 3**, line 39) and integrated into the existing POA graph (**Algorithm 3**, line 40).

**Figure 4.**
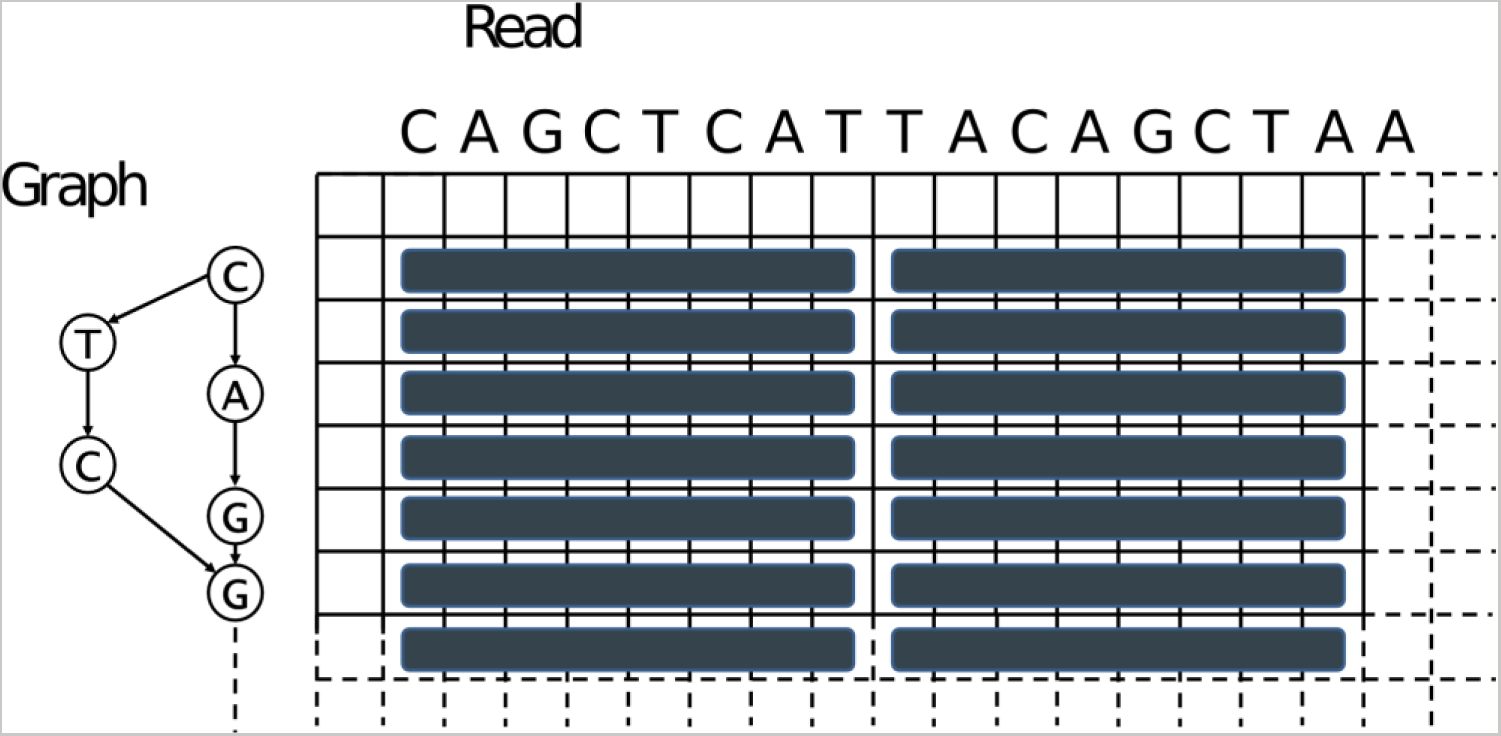
Depiction of the SIMD vectorization approach used in SPOA.

SIMD intrinsics decrease the time complexity for alignment from *O*((2*n_p_* + 1) *n*|*V*|) to roughly *O*((2*n_p_*/*k* + 1) *n*|*V*|), where *k* is the number of variables that fit in a SIMD vector, *n_p_* is the average number of predecessors in the graph and |*V*| is the number of nodes in the graph. SPOA supports Intel SSE version 4.1 and higher, which embed 128 bit registers. Both short (16 bits) and long (32 bits) integer precisions are supported (therefore *k* equals 8 and 4 variables, respectively). 8 bit precision is insufficient for the intended application of SPOA and is therefore not used. Alongside global alignment displayed in **Algorithm 3.**, SPOA supports local and semi-global alignment modes, in which SIMD vectorization is implemented as well.

**Algorithm 3.**
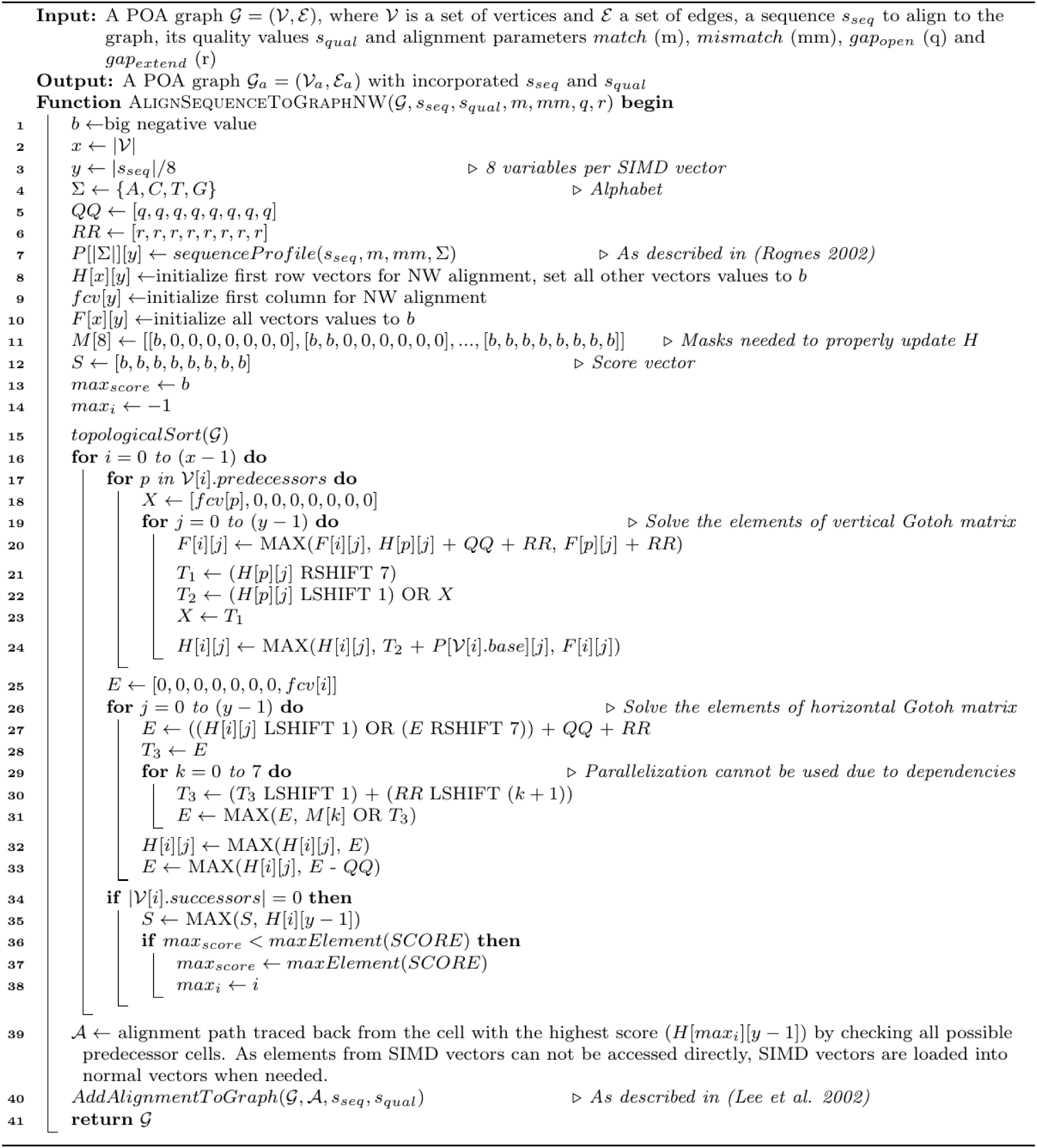
Pseudocode for the SPOA algorithm. The displayed function aligns a sequence to a pre-constructed POA graph using SIMD intrinsics. Capitalized variables are SIMD vectors. Alignment mode is Needleman-Wunsch.

### Implementation and reproducibility

Racon and SPOA are both implemented in C++. All tests were run using Ubuntu based systems with two 6-core Intel(R) Xeon(R) E5645 CPUs @ 2.40GHz with Hyperthreading, using 12 threads where possible. The versions of various methods used in the comparisons reported here are:

- Minimap - https://github.com/lh3/minimap.git, commit: 1*cd*6*ae*3*bc*7*c*7
- Miniasm - https://github.com/lh3/miniasm.git, commit: 17*d*5*bd*12290*e*
- Canu - https://github.com/marbl/canu.git, version 1.2, commit: *ab*50*ba*3*c*0*cf*0.
- FALCON-integrate project - https://github.com/PacificBiosciences/FALCON-integrate.git, commit: 8*bb*2737*fd*1*d*7.
- Nanocorrect - https://github.com/jts/nanocorrect.git, commit: *b*09*e*93772*ab*4
- Nanopolish - https://github.com/jts/nanopolish.git, commit: 47*dcd*7*f*147*c*.
- MUMmer - DNAdiff version 1.3, NUCmer version 3.1.

### Datasets

Five publicly available PacBio and Oxford Nanopore datasets were used for evaluation. These are:

1. *Lambda* phage, Oxford Nanopore, ENA submission ERA476754, with 113× coverage of the NC_001416 reference genome (48502bp). Link: ftp://ftp.sra.ebi.ac.uk/vol1/ERA476/ERA476754/oxfordnanopore_native/Lambda_run_d.t_ar.gz. This dataset was subsampled to coverages of 30× and 81× for testing.
2. *E. coli* K-12 MG1655 SQK-MAP006-1 dataset, Oxford Nanopore, R7.3 chemistry, 54× pass 2D coverage of the genome (U00096.3, 4.6Mbp). Link: http://lab.loman.net/2015/09/24/first-sqk-map-006-experiment/
3. *E. coli* K-12 PacBio P6C4 PacBio dataset, 160× coverage of the genome (U00096.3). The dataset was generated using one SMRT Cell of data gathered with a PacBio RS II System and P6-C4 chemistry on a size selected 20kbp library of *E. coli*K-12. Link: https://s3.amazonaws.com/files.pacb.com/datasets/secondary-analysis/e-coli-k12-P6C4/p6c4_ecoli_RSII_DDR2_with_15kb_cut_E01_1.tar.gz
4. *S. cerevisiae* W303 P4C2 PacBio dataset (https://github.com/PacificBiosciences/DevNet/wiki/Saccharomyces-cerevisiae-W303-Assembly-Contigs). The dataset is composed of 11 SMRT cells, of which one was not used in this study because the containing folder (“0019”) was incomplete and the data could not be extracted. The S288C reference (12.1 Mbp) was used for comparison (http://downloads.yeastgenome.org/sequence/S288C_reference/chromosomes/fasta/). Coverage of the dataset with respect to the S288C reference is approx. 127×.
5. *C. elegans*, a Bristol mutant strain, 81× coverage of the genome (gi|449020133). The dataset was generated using 11 SMRT cells P6-C4 chemistry on a size selected 20kbp library. Link: https://github.com/PacificBiosciences/DevNet/wiki/C.-elegans-data-set

### Evaluation methods

The quality of called consensus sequences was evaluated primarily using Dnadiff (Delcher et al. 2003). The parameters we took into consideration for comparison include: total number of bases in the query, aligned bases on the reference, aligned bases on the query and average identity. In addition, we measured the time required to perform the entire assembly process by each pipeline.

The quality of error-corrected reads was evaluated by aligning them to the reference genome using GraphMap (Sović et al. 2016b) with settings “-a anchorgotoh”, and counting the match, mismatch, insertion and deletion operations in the resulting alignments.

## Data access

No new sequencing datasets were generated in the course of this study.

## Acknowledgements

This work has been supported in part by Croatian Science Foundation under the project UIP-11-2013-7353. IS is supported in part by the Croatian Academy of Sciences and Arts under the project “Methods for alignment and assembly of DNA sequences using nanopore sequencing data”. NN is supported by funding from A*STAR, Singapore

## Author contributions

IS, MS and RV devised the original concept for Racon. IS developed and implemented several prototypes for Racon. RV suggested and implemented the SIMD POA version. IS implemented a version of Racon which uses SPOA for consensus generation and error-correction. IS and RV tested Racon and SPOA. MS and NN provided helpful discussions and guidance for the project. MS helped define applications and devised tests for the paper; IS ran the tests, and collected and formatted the results. IS and RV wrote the paper with input from all authors. MS and RV designed the figures for the paper. NN proof-read and corrected parts of the paper. All authors read the paper and approve of the final version. MS coordinated the project and provided computational resources.

## Code Availability

Racon and SPOA are available open source under the MIT license at https://github.com/isovic/racon.git and https://github.com/rvaser/spoa.git.

